## Supplemental Text, Tables and Figures for "The Impact of Antimalarial Resistance on the Genetic Structure of *Plasmodium falciparum* in the DRC"

### TABLE OF CONTENTS


1. SUPPLEMENTAL METHODS Page 3
2. Supplemental Table 3 Page 7
3. Supplemental Figure 1 Page 8
4. Supplemental Figure 2 Page 9
5. Supplemental Figure 3 Page 10
6. Supplemental Figure 4 Page 11
7. Supplemental Figure 5 Page 12
8. Supplemental Figure 6 Page 13
9. Supplemental Figure 7 Page 14
10. Supplemental Figure 8 Page 15
11. Supplemental Figure 9 Page 16
12. Supplemental Figure 10 Page 17
13. Supplemental Figure 11 Page 18
14. Supplemental Figure 12 Page 19
15. Supplemental Figure 13 Page 20
16. SUPPLEMENTAL REFERENCES Page 21

### SUPPLEMENTAL METHODS

**MIP Design:** We used two distinct MIP panels - a genome-wide panel designed to capture overall levels of differentiation and relatedness, and a drug resistance panel aimed at polymorphic sites known to be associated with antimalarial resistance. The drug resistance MIP panel was described previously[1]. When selecting targets for the genome-wide panel, we used publicly available *Plasmodium falciparum* whole genome sequences provided by the Pf3k project (Data Release 5) and *P. falciparum* Community project (Data Release 4), which are part of the wider MalariaGEN Consortium[2,32]. This consisted of 923 samples in total, from Cameroon (n=134), the Democratic Republic of the Congo (n=285), Kenya (n=52), Malawi (n=369), Nigeria (n=5), Tanzania (n=66), and Uganda (n=12) (**Supplemental Table 2**). The genomic sequence from these samples underwent alignment, variant calling, and variant-filtering following the Pf3k strategy consistent with the Genome Analysis Toolkit (GATK) Best Practices with minor modifications[3–6]. Reads were aligned to the *P. falciparum* 3d7 reference assembly genome (version 3) using BWA-MEM with a raised base-match bonus (A=2) and clip penalty for local alignment (L=15) for increased sensitivity and specificity through hypervariable regions, and with all other flags set to default[7,8]. Given that there were paired-end reads, we used samtools fixmates, to synchronize any overlapping paired-end bases[9]. Mate-fixed reads were then deduplicated and merged with Picard Tools MarkDuplicates, and MergeSamFile (version 2.2.4), respectively[10]. Finally, we performed local realignment of complex regions using Genome Analysis Toolkit (GATK) IndelRealigner (version 3.6).

Following best practices for variant calling, we first used the GATK BaseRecalibrator tool to adjust our samples' base-quality scores using the the sequences from the *P. falciparum* Genetic Cross project as the training set. Variant discovery was performed separately on each recalibrated binary alignment map (BAM) file using GATK HaplotypeCaller with the minimum Phred score for a variant to be called at 30 and a ploidy of one. Setting the ploidy to one shifts the genotype-call to the major haplotype in polyclonal infections, in expectation. The individual variant call files (VCFs) then underwent joint variant discovery with the GATK GenotypeGVCFs tool. Following this step, we used the GATK VariantRecalibrator and ApplyRecalibration tools to recalibrate our discovered single-nucleotide-polymorphisms (SNPs) and insertion-deletions (INDELs). For SNP recalibration tuning, we again used the *P. falciparum* genetic crosses as the training set with the quality depth, mapping quality, fisher-score, strand-odds ratio, and allele depth as the covariate considered and the maximum gaussian clusters set to eight processes. Similarly, for INDEL recalibration, the *P. falciparum* cross data was again used to train the model but only quality depth, fisher-score, strand-odds ratio, and allele depth were considered as covariates and the maximum gaussian clusters were set to four processes. In the ApplyRecalibration step, the truth sensitivity level for filtering was set at 99% for both SNPs and INDELs. Finally, from the recalibrated-joint VCF, we filtered all variants with a variant-quality recalibration log-odds of less-than or equal to zero. In addition, variants were excluded if they were not within the "core" genome as defined by the Pf3k project.

To further decrease our false discovery rate, we subsetted to biallelic SNPs and excluded all samples (original BAMs) with fewer than 70% of loci that were callable as determined by GATK CallableLoci with flags set to a minimum base quality of 20, minimum mapping quality of 10, and a minimum depth of 4. In addition, we excluded samples from Uganda and Nigeria, as these countries did not have enough high-quality genomes for analysis. Next, we separated the joint-VCF into country-level VCF and excluded intervals that had fewer than 5-fold coverage at 50% of loci within a given country using GATK CoveredByNSamplesSites (version 3.4.46). The country-level VCFs were then re-merged and annotated using snpEFF and the Pf3D7v91 genome pre-package in the snpEFF databases.

From this filtered-VCF, we calculated Weir and Cochram's $F_{ST}$ with respect to country for each biallelic locus[11]. The 1,000 loci with the highest $F_{ST}$ values were considered for MIP design as phylogeographically informative loci. Of these 1,000 potential loci, 739 were identified as regions that were suitable for MIP-probe design. Separately, from the combined SNP file, we identified 1,595 potential loci that had a minor-allele frequency greater than 5%, had an $F_{ST}$ value between 0.005 and 0.2, and were annotated by SNPEff as functionally silent mutations. These were identified as putatively neutral SNPs. Of these 1,595 potential loci, 1151 were suitable for MIP-probe design. 76 loci were shared between phylogeographically informative and putatively neutral loci.

**Complexity of Infection:** We applied THE REAL McCOIL categorical method to the SNP genotyped samples to estimate each individual's COI[12], using a 10% minor allele frequency cutoff for calling a heterozygous locus. We performed five repetitions for each sample, with a burn-in period of $10^4$ iterations followed by $10^6$ sampling iterations and using standard methodology to confirm convergence between chains[13]. Given the concurrent estimation of population allele frequencies within THE REAL McCOIL, samples were grouped initially within their countries. Default priors were assigned for each parameter, with a maximum observable COI equal to 25 and sequencing measurement error estimated along with COI and allele frequencies. COI estimates were compared between countries using 100,000 repetitions of a non-parametric bootstrap to estimate the 95% confidence interval from the bootstrapped COI density. Additionally, to test for a relationship between COI and transmission intensity, we modelled the exponential relationship between COI and malaria prevalence in the DRC at the cluster level, with a random intercept for each administrative region.

**Extended haplotype homozygosity analysis:** Alleles at these biallelic SNPs were polarized as ancestral versus derived using P. reichenowi as an outgroup. Briefly, the *P. falciparum* 3D7 assembly was aligned to the PlasmoDB v38 assembly of the *P. reichenowi* CDC strain with nucmer[14] using parameters "-g 500 -c 500 -l 10" as in Otto et al.[15]. Only segments with globally unique, one-to-one alignments were retained. The *P. reichenowi* allele was defined as ancestral at all SNPs in these segments; SNPs falling outside these segments were considered to have ambiguous ancestral state and were excluded from analysis. In order to account for linkage between SNPs, we created a recombination map from the pedigrees reported in Miles et al.[16] under the assumption of no unobserved double-crossovers. The genetic positions of SNPs in the MIP panel were interpolated on this map with piecewise-linear interpolation.

We then subsetted to a set of monoclonal samples as identified by THE REAL McCOIL categorical method. Haplotypes were created from the genotype calls, which were the majority within-sample allele frequency. To account for population structure, we analyzed the samples with respect to country with the exception of the DRC, which was split into two groups using K-means clustering weighted by longitude- and latitude-coordinates (**Supplementary Figure 7**). K-means clustering with two groups resulted in an East-West divide that is consistent with previous publications of DRC population substructuring[17]**.** Samples were further subsetted to those without any missing genotype data.

Given that the MIP panel density was not uniform or symmetric by design, we did not perform genome-wide scans for recent positive selection[18,19]. Genome-wide scans for recent positive selection with our MIP panel would have been biased towards sites with higher MIP density and would not have been comparable across the genome. However, given that MIP-site densities are identical between subpopulations at the same genomic region, cross-population interpretations of differing selection pressures were still considered valid. As a result, we calculated the log-ratio of the integrated EHH for the derived allele, hereafter called the $XP\text{-}EHH_D$, to differentiate recent positive selection between subpopulations[20]. We focused on sites that were identified in this study as putative drug-resistance loci and had a prevalence of at least 10% in the DRC. The resulting $XP\text{-}EHH_D$ were then standardized and a one-sided p-value (p'XP-EHH) was calculated for each site assuming a Gaussian cumulative distribution[21,22].

### SUPPLEMENTAL TABLES

#### Supplemental Table 3 - XP-EHH statistics

| region.x | name | mut_name | marker | region.y | crudexpehh | scaledxpehh | pprimexpehh_scale | statsig |
|---|---|---|---|---|---|---|---|---|
| DRC-East | dhfr | N51I | 8 | DRC-West | 0.193192477 | 0.619466291 | 0.482417688 | FALSE |
| DRC-East | dhfr | C59R | 9 | DRC-West | -0.253243174 | 0.102909101 | 0.401389585 | FALSE |
| DRC-East | dhfr | S108N | 10 | DRC-West | -0.109349489 | 0.269404133 | 0.414850174 | FALSE |
| DRC-East | mdr1 | N86Y | 34 | DRC-West | NA | NA | NA | NA |
| DRC-East | mdr1 | Y184F | 39 | DRC-West | -0.075231395 | 0.308881147 | 0.419807423 | FALSE |
| DRC-East | mdr1 | D1246Y | 47 | DRC-West | NA | NA | NA | NA |
| DRC-East | crt | M74I | 16 | DRC-West | -1.777313372 | -1.660546432 | 0.997854875 | TRUE |
| DRC-East | crt | N75E | 17 | DRC-West | -1.777313372 | -1.660546432 | 0.997854875 | TRUE |
| DRC-East | crt | K76T | 19 | DRC-West | -1.777313372 | -1.660546432 | 0.997854875 | TRUE |
| DRC-East | crt | I356T | 25 | DRC-West | NA | NA | NA | NA |
| DRC-East | dhps | S436A | 37 | DRC-West | 0.42714805 | 0.890169209 | 0.571157673 | FALSE |
| DRC-East | dhps | G437A | 38 | DRC-West | 0.005924775 | 0.402784497 | 0.434318893 | FALSE |
| DRC-East | dhps | K540E | 39 | DRC-West | 0.696155866 | 1.201430017 | 0.712527663 | FALSE |
| DRC-East | dhps | A581G | 40 | DRC-West | 0.317580348 | 0.763391735 | 0.525636168 | FALSE |
| DRC-East | mdr2 | I492V | 5 | DRC-West | -0.34771804 | -0.006404909 | 0.399098842 | FALSE |
| DRC-East | mdr2 | F423Y | 6 | DRC-West | 0.02910711 | 0.429608076 | 0.439167302 | FALSE |

### SUPPLEMENTAL FIGURES

**Supplemental Figure 1 - UMI depth distributions**. Histograms show the raw distribution of coverage (number of unique UMIs) per locus for the genome-wide and drug resistance MIP panels on a log scale.

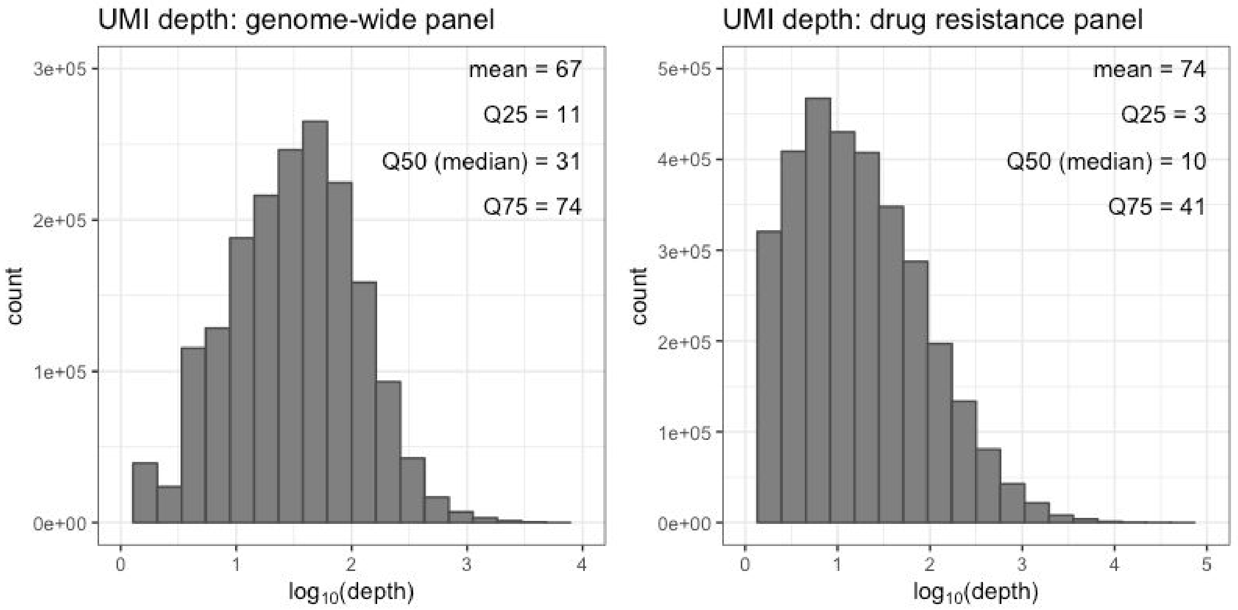

**Supplemental Figure 2 - Expected vs. measured allele frequencies in control samples for genome-wide MIPs.** A mix of 4 laboratory strains (as described in the methods) were used as controls. Each targeted SNP's expected allele frequency (in increasing order) is plotted in blue based on which strains harbor the SNP and what ratio the strain was mixed in the sample. Each SNP's frequency as measured experimentally is plotted in red. Pearson's correlation coefficient between the expected and observed frequencies were 0.968 ($R^2$=0.938). In total, 114 control reactions were run with experiments.

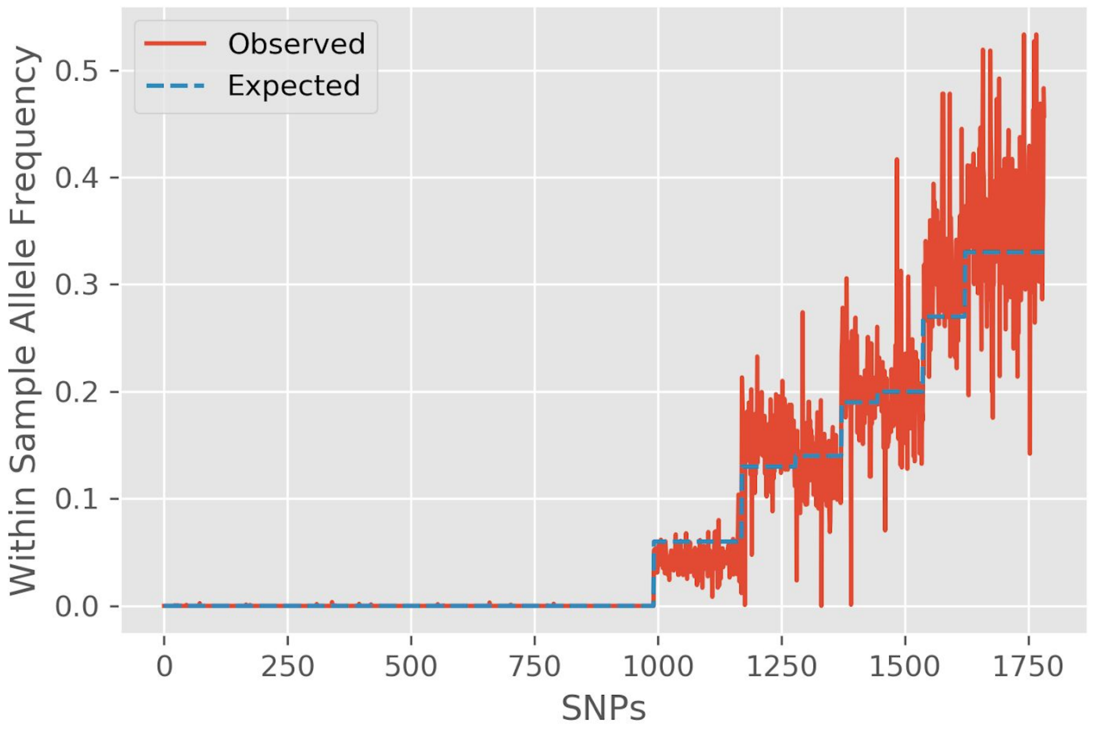

**Supplemental Figure 3 - COI distribution per country.** Violin plots show the distribution of the estimated COI, which is shown as the median from the posterior distribution estimated using THE REAL McCOIL categorical method. The mean and 95% confidence interval, estimated using a non-parametric bootstrap, are shown in red. Samples collected from Ghana had a significantly lower COI compared to the other countries, and samples collected from Zambia had a significantly higher COI compared to DRC.

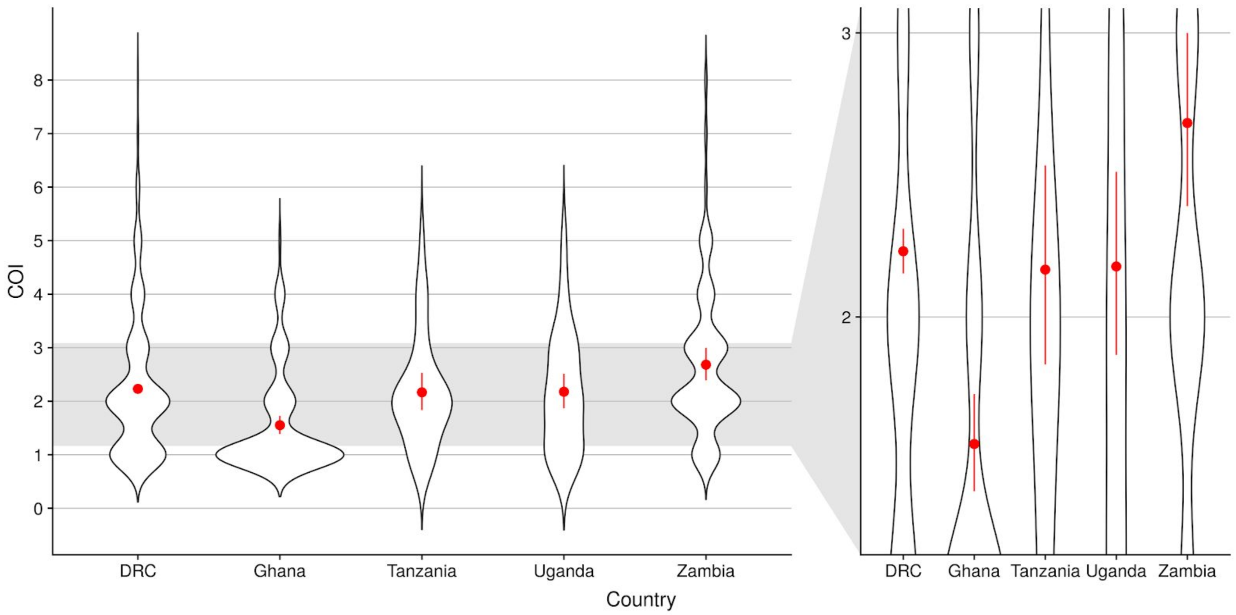

**Supplemental Figure 4 - Relationship between COI and prevalence.** In **(a)** the relationship between COI  and microscopy prevalence at the province level is shown for the samples collected from the DRC. Each point represents the survey-weighted COI estimate and a locally weighted regression is shown in blue with the 95% confidence interval shaded in grey. The same relationship at the cluster level is shown in **(b)** with the size of the points reflecting the survey-weighted sample size.

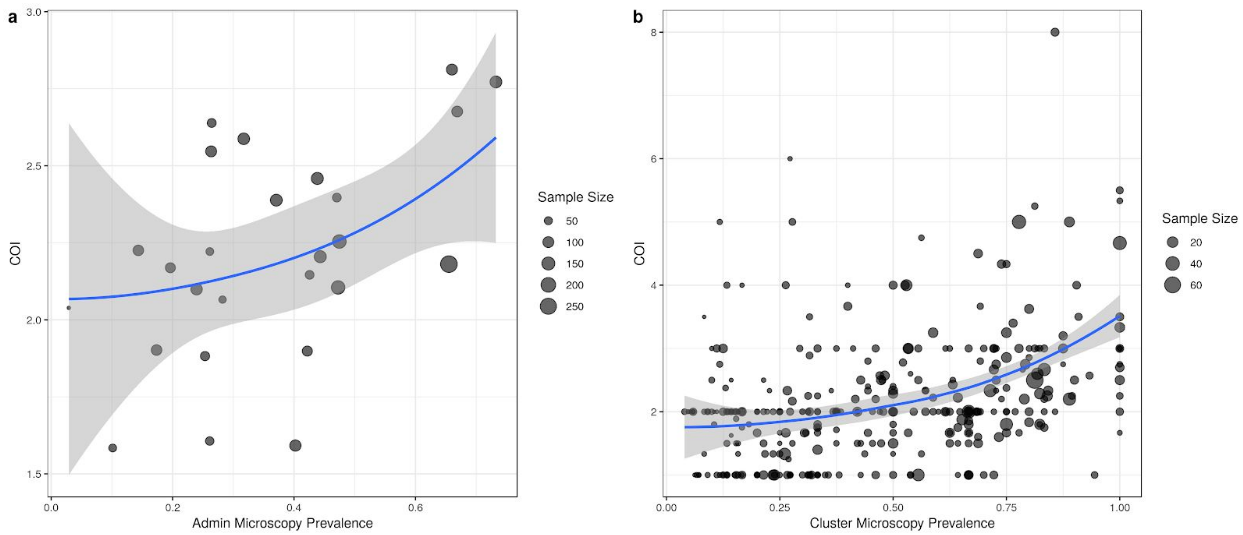

**Supplemental Figure 5 - PCA variance explained**. The variance explained by the first 20 principal components as a percentage of overall variance.

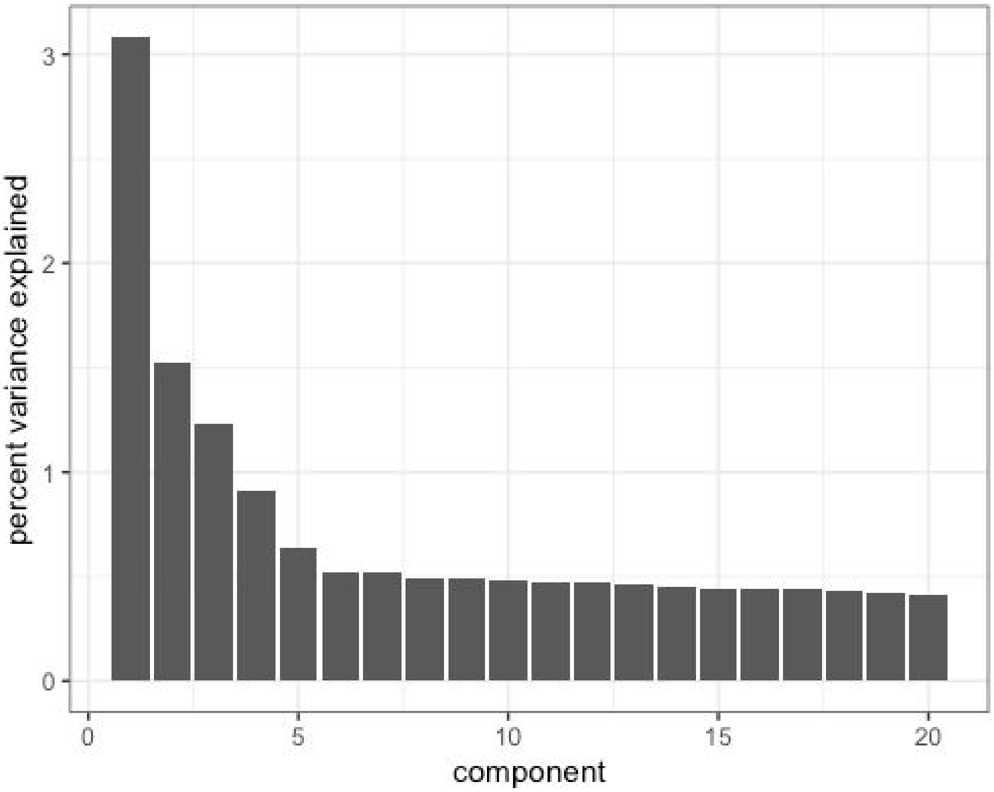

**Supplemental Figure 6 - Within-sample allele frequencies of highly related samples**. For the 12 sample pairs identified as highly related (IBD>0.9), scatterplots compare the raw within-sample allele frequencies (WSAF) at every locus. Perfectly matching monoclonal genotypes would be represented as a single point in the lower-left and upper-right corners, however, polyclonal infections and sequencing errors cause deviations from this pattern. The number of loci that match or mismatch in terms of occupying the same or different intervals in {[0,0.5), [0.5,1]} is shown.

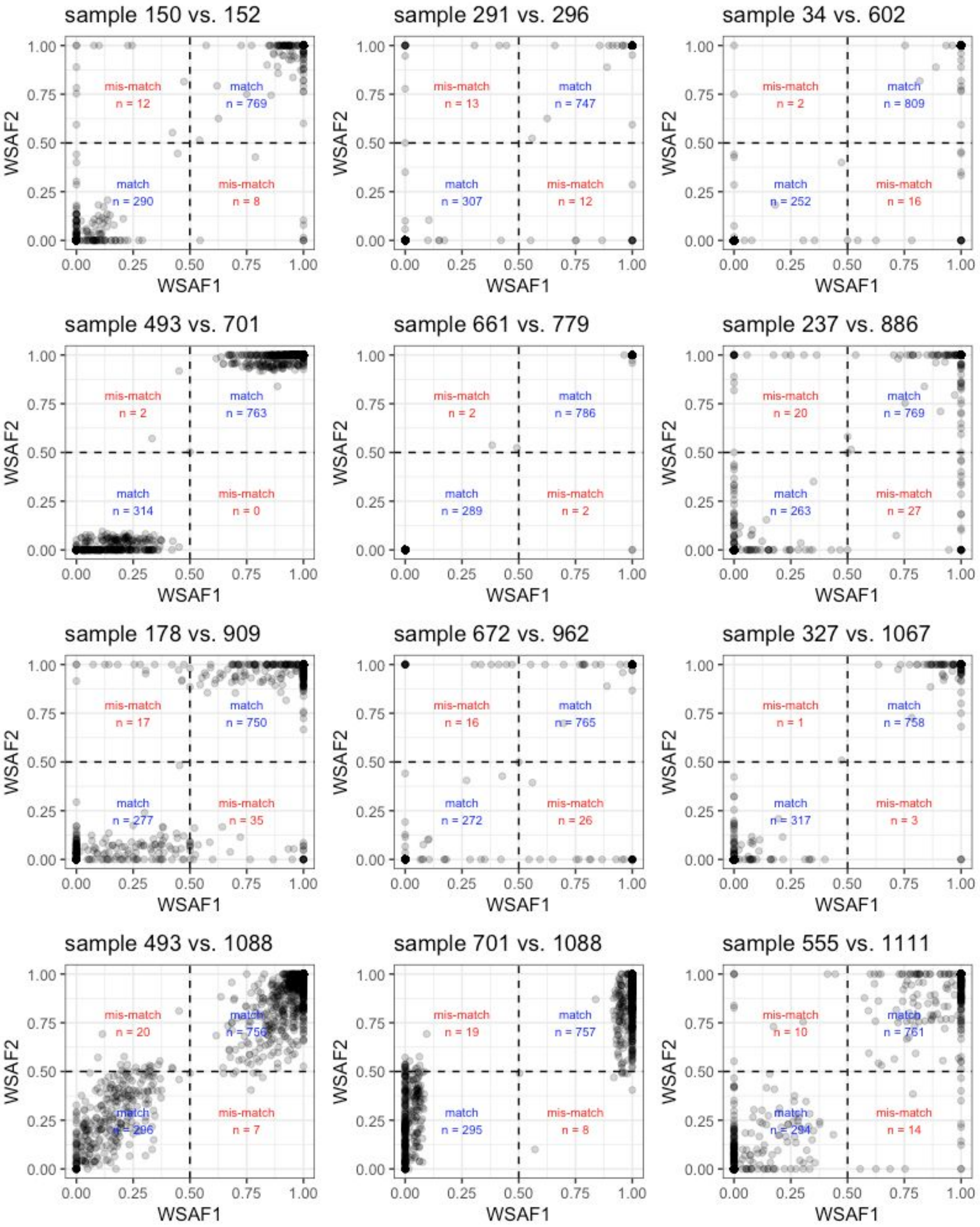

**Supplemental Figure 7 - Population K-means clustering**. Geographic distribution of samples in each of the two clusters produced by K-means clustering within DRC. Countries outside DRC remain single groups.

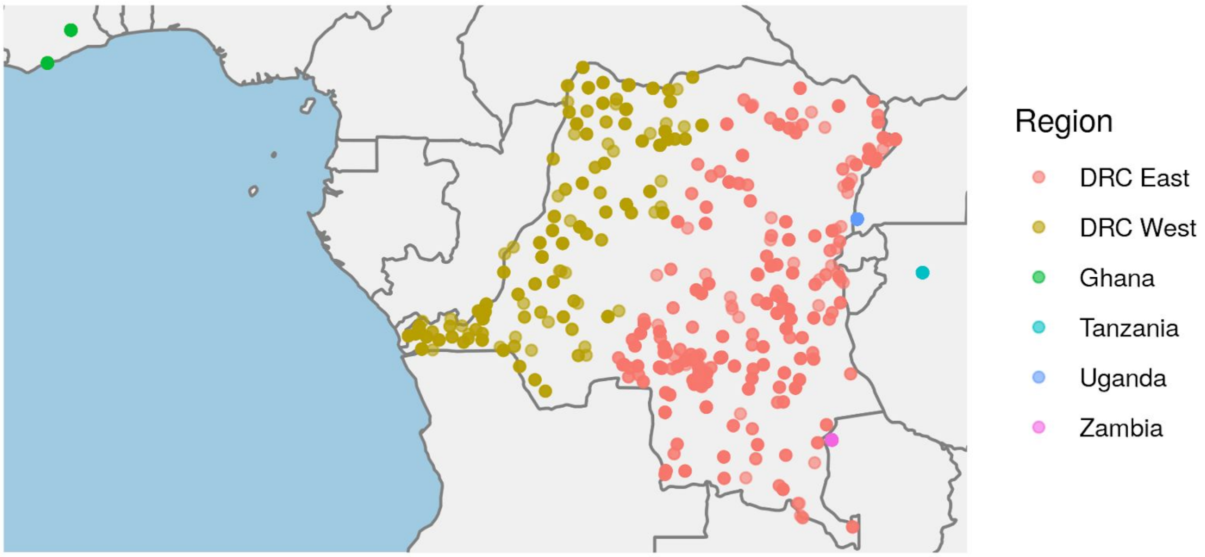

**Supplemental Figure 8 - *pfcrt* I356T EHH and haplotype plots among monoclonal infections with no missing genotype data**. Panels (a) and (b) show the EHH decay plots 200 kilobases upstream and downstream of the I356T core SNP in centimograns respect to the eastern DRC and western DRC, respectively. Panels (c) and (d) display the extended haplotypes with the SNPs colored at each respective loci contributing to the ancestral and derived extended haplotype. The core SNP is marked with a black asterisk.

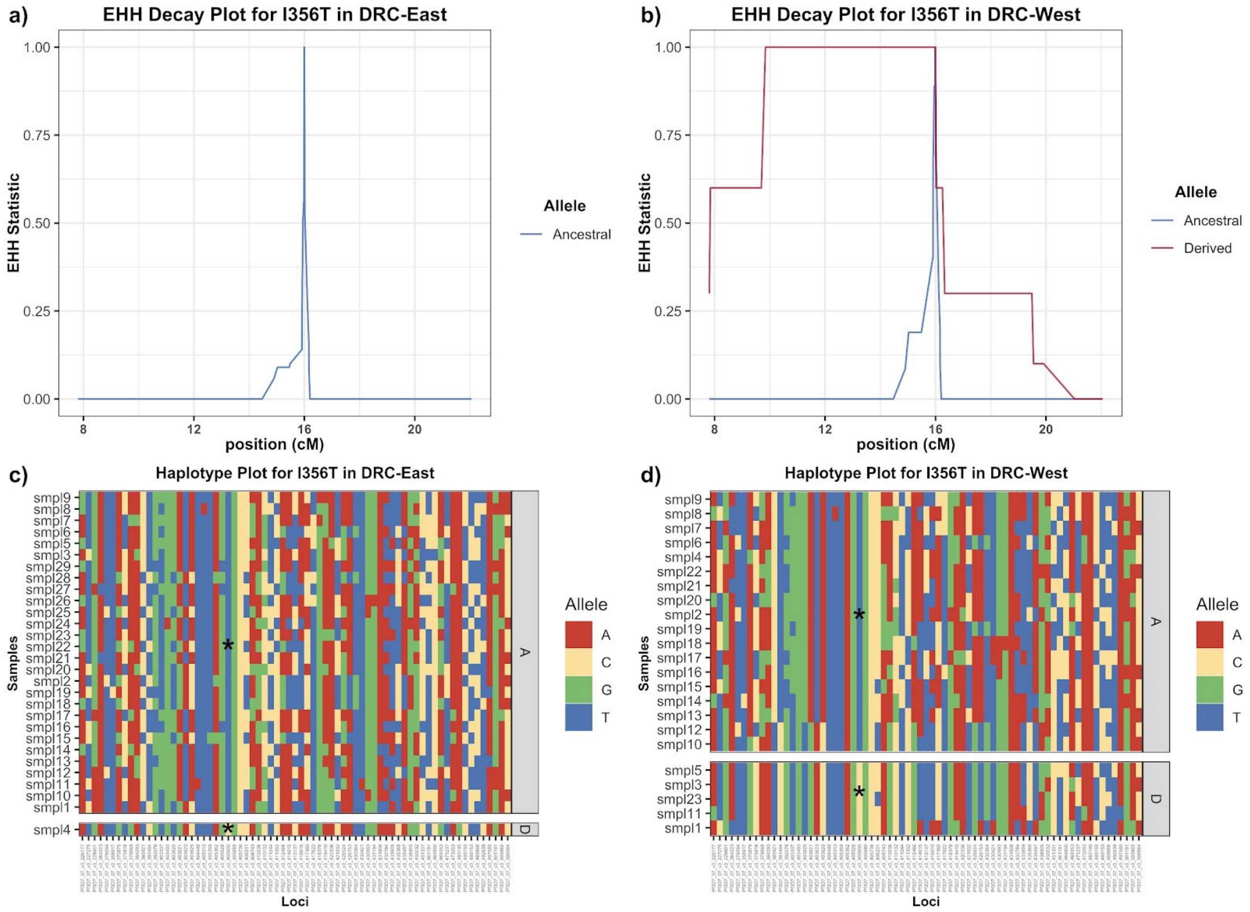

**Supplementary Figure 9 - *dhps* A581G EHH and haplotype plots among monoclonal infections with no missing genotype data**. Panels (a) and (b) show the EHH decay curve 200 kilobases upstream and downstream of the A581G core SNP in centimograns respect to the eastern DRC and western DRC, respectively. Panels (c) and (d) display the extended haplotypes with the SNPs colored at each respective loci contributing to the ancestral and derived extended haplotype. The core SNP is marked with a black asterisk.

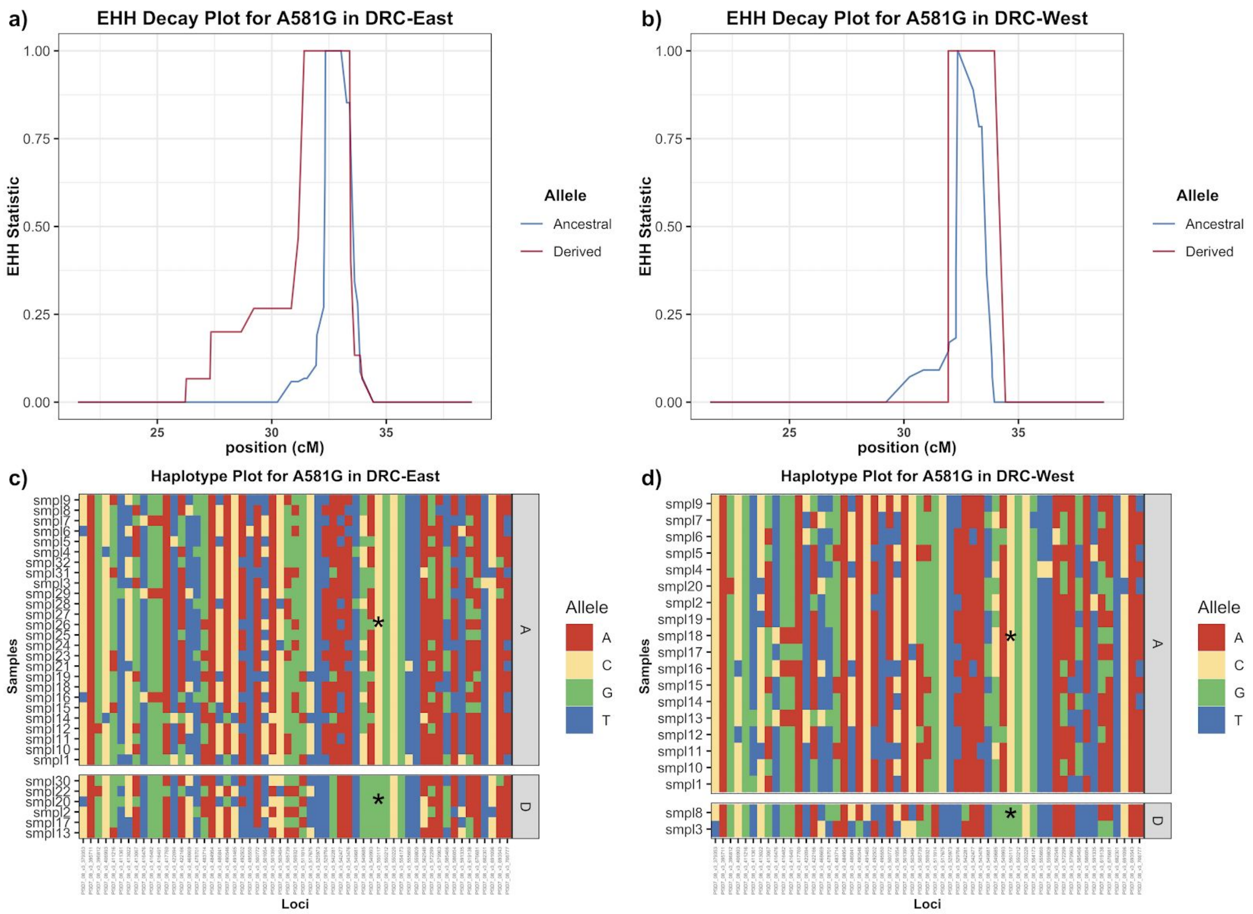

**Supplementary Figure 10 - *dhps* K540E EHH and haplotype plots among monoclonal infections with no missing genotype data**. Panels (a) and (b) show the EHH decay curve 200 kilobases upstream and downstream of the K540E core SNP in centimograns respect to the eastern DRC and western DRC, respectively. Panels (c) and (d) display the extended haplotypes with the SNPs colored at each respective loci contributing to the ancestral and derived extended haplotype. The core SNP is marked with a black asterisk.

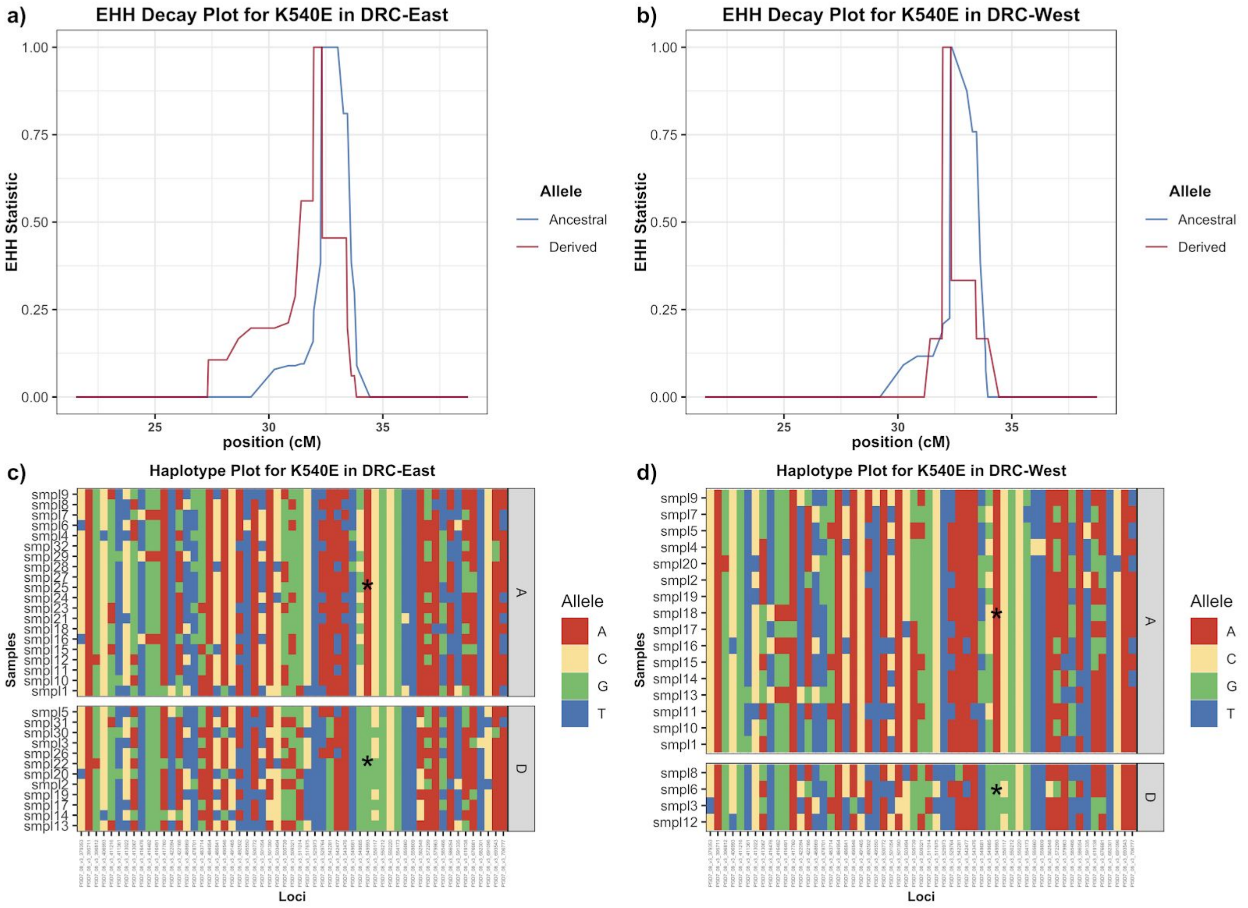

**Supplementary Figure 11 - *dhps* G437A EHH and haplotype plots among monoclonal infections with no missing genotype data**. Panels (a) and (b) show the EHH decay curve 200 kilobases upstream and downstream of the G437A core SNP in centimograns respect to the eastern DRC and western DRC, respectively. Panels (c) and (d) display the extended haplotypes with the SNPs colored at each respective loci contributing to the ancestral and derived extended haplotype. The core SNP is marked with a black asterisk.

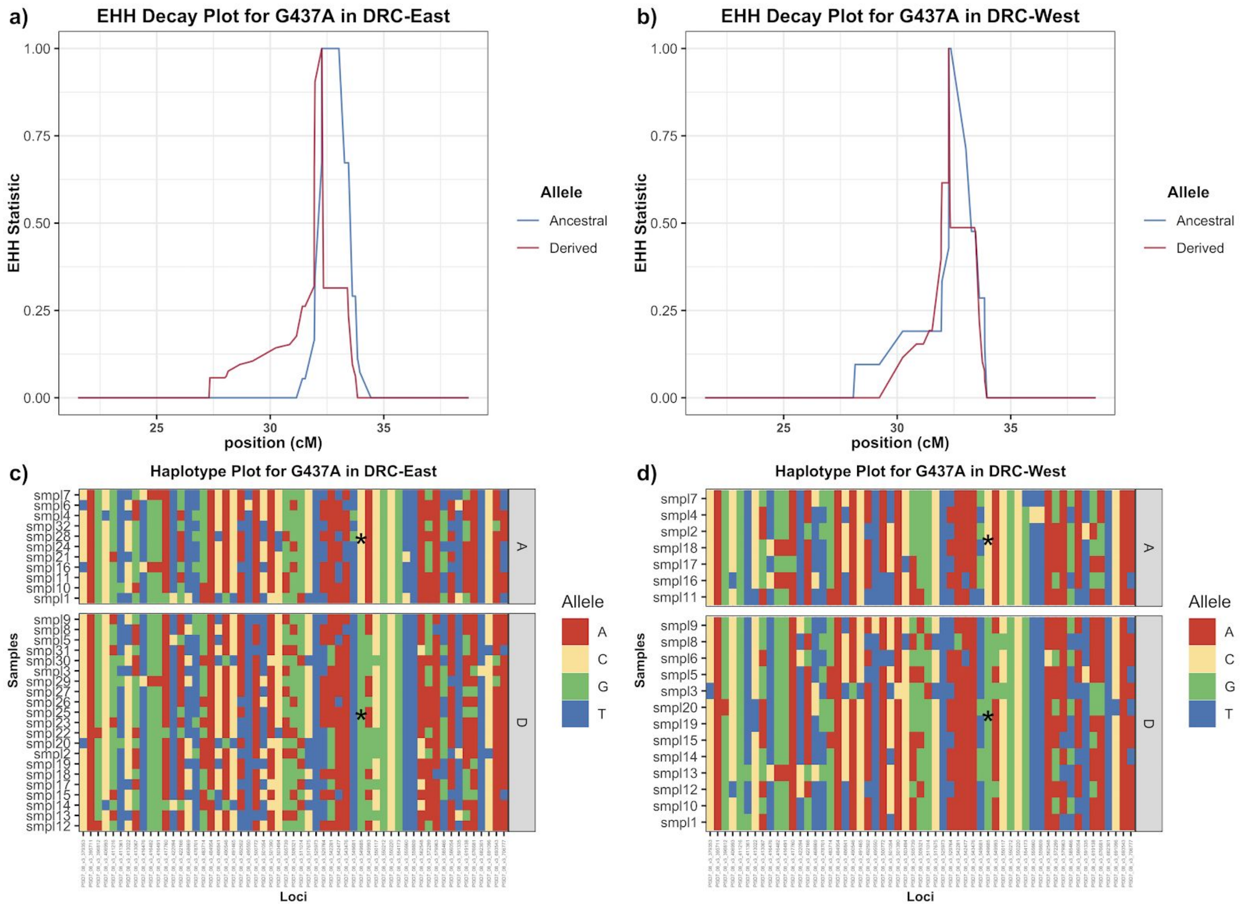

**Supplementary Figure 12 - Remaining EHH decay plot for the *mdr1*, *mdr2*, and *dhfr* genes among monoclonal infections with no missing genotype data**. Plots (a-j) display the EHH decay curve 200 kilobases upstream and downstream of the respective core SNP. The remaining loci did not appear to be under recent positive selection.

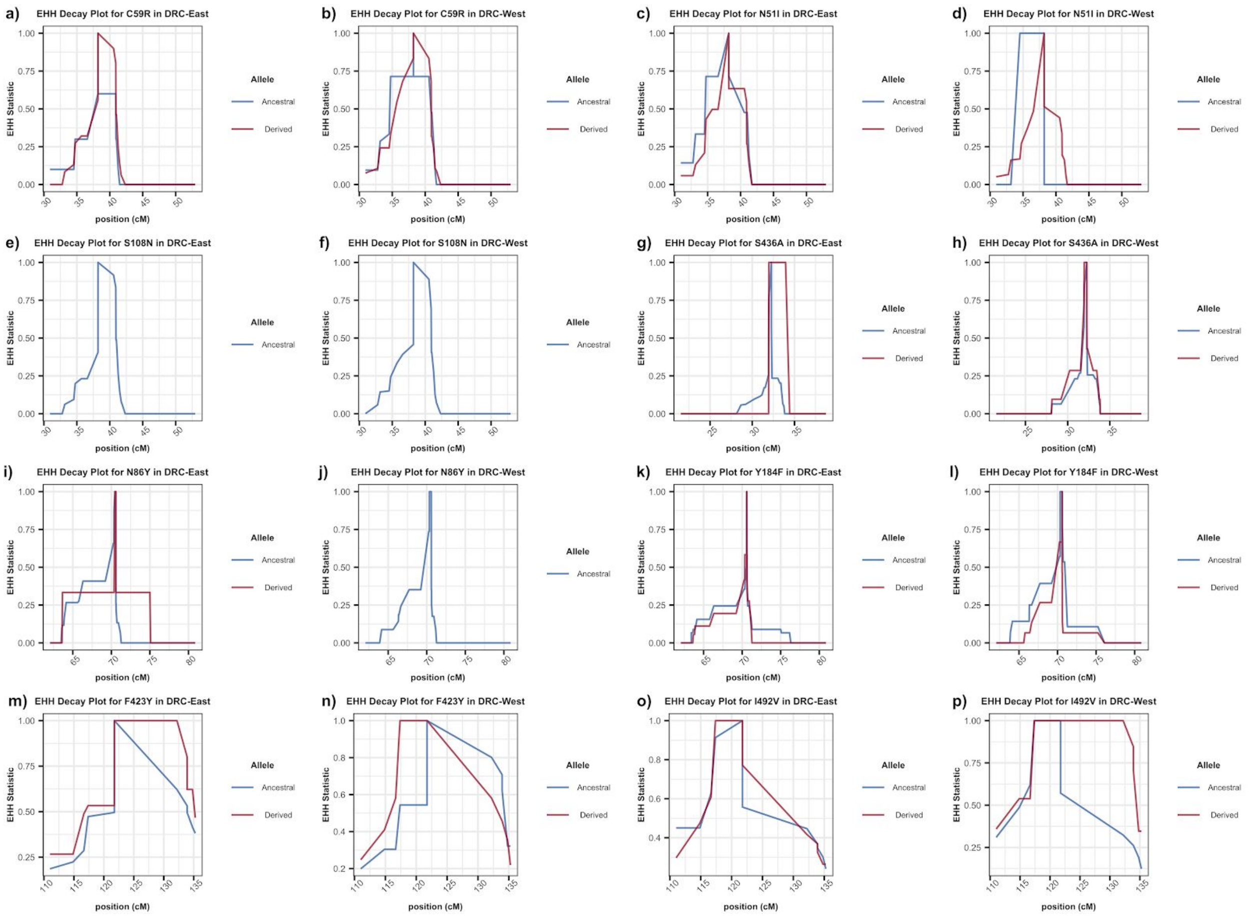

**Supplemental Figure 13 - Genomic positions of MIP targets.** Genomic locations of geographically informative (red) and putatively neutral (blue) SNP targets are indicated on each chromosome.

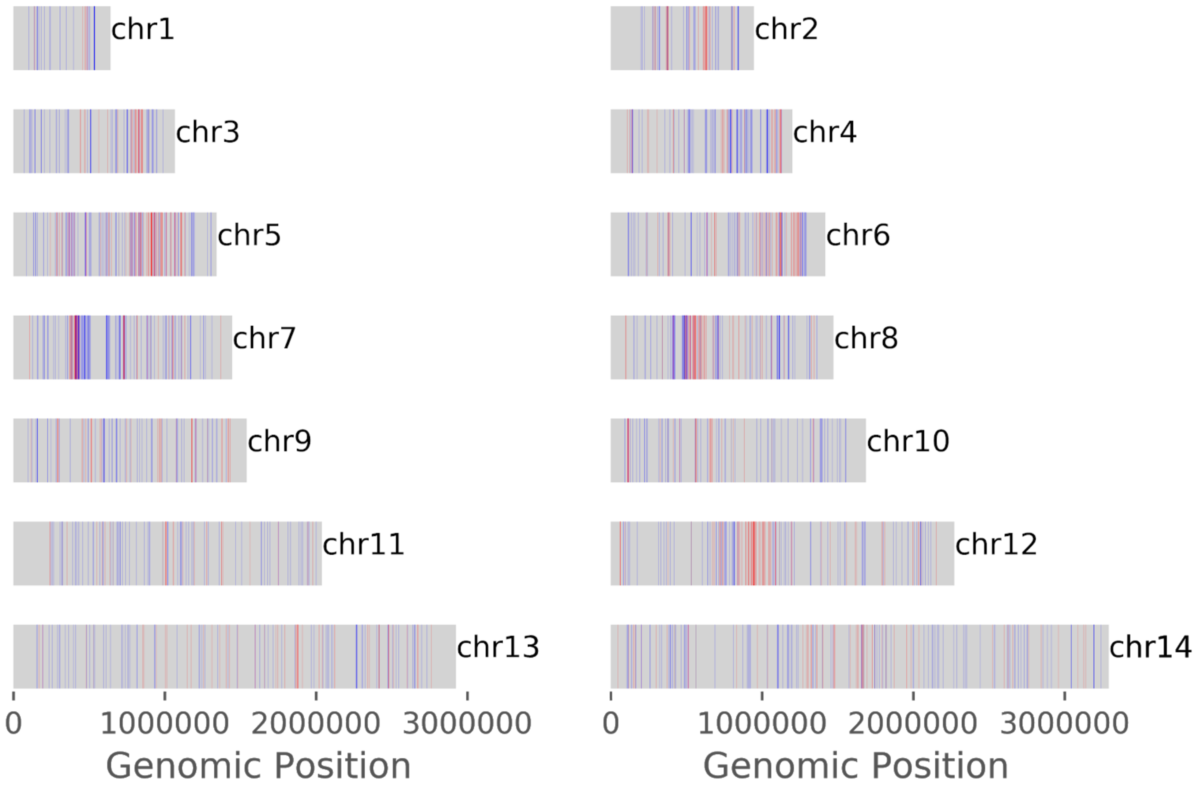
